# GeneInsight: Condensing Gene Set Knowledge via Language Models

**DOI:** 10.1101/2025.07.07.663611

**Authors:** Wee Loong Chin, Kevin Chen, Timo Lassmann

## Abstract

Interpreting gene sets is often complicated by the overwhelming number of annotations associated with individual genes, making it difficult to extract meaningful biological insights. To address this issue, we developed GeneInsight, an AI-powered tool that combines advanced topic modelling with large language models to automatically synthesise diverse biological annotations from literature, gene ontologies, and databases such as STRING. GeneInsight consolidates extensive annotations into coherent thematic summaries that render such data readily interpretable, thereby enabling the rapid extraction of biologically significant insights that conventional enrichment analyses often overlook.

## Main text

Gene set interpretation is a fundamental task in functional genomics, where researchers must derive biological insights from lists of genes identified in high-throughput experiments. Current approaches utilise statistical enrichment methods that query predefined functional databases, such as Gene Ontology^1^ and KEGG pathways^2^, to identify overrepresented biological processes^3,4^. Although powerful, current methods yield fragmented outputs, such as lists of enriched terms from various ontologies, leaving researchers to manually integrate these results to achieve functional insights, a process that is both inefficient and error-prone.

Exploratory gene set analysis has become increasingly challenging as the volume of annotated datasets grows. Researchers compare their findings not only with gene ontology term enrichments but also with signatures from knockdown experiments and diverse resources such as LINCS (Library of Integrated Network-based Cellular Signatures)^5^ and the STRING database (STRING-DB)^6^. The main issue is that enrichment analysis tests for the over-representation of genes associated with specific terms. When gene lists overlap, the same genes often appear under many different functional terms. As a result, enrichment analysis can return a large number of distinct-sounding terms that all point to the same underlying biology, making interpretation more difficult.

However, the challenge extends beyond simply removing duplicate information — simplistic filtering of overlapping gene sets would obscure important biological relationships that only become apparent when analysing genes across the diverse resources mentioned above. These relationships often represent biological processes that bridge multiple databases and reveal insights not captured by any single resource. Consequently, manually curating enrichment outputs to identify both redundancies and meaningful biological patterns is not only error-prone and prohibitively time-consuming but also risks overlooking crucial biological connections.

We hypothesise that recently introduced topic modelling techniques can address this problem. By analysing how terms co-occur across texts, topic modelling reveals underlying themes without requiring predefined categories. These unsupervised statistical methods – including Latent Dirichlet Allocation (LDA)^7^ and Non-negative Matrix Factorization (NMF)^8^ – identify recurring patterns in document collections. The resulting latent themes represent collections of related terms that frequently appear together and may correspond to biological processes, pathways, or functional modules not explicitly defined in current annotation databases.

Recent advances in large language models^9^ (LLMs) and topic modelling create powerful new opportunities for gene set interpretation. LLMs have demonstrated remarkable capabilities in contextual understanding and natural language generation, potentially enabling automated synthesis of distributed biological knowledge. Here we present GeneInsight, a tool that integrates LLMs with topic modelling to automate gene set interpretation^10^. Our system aggregates gene-specific annotations from the STRING database, applies topic modelling to identify coherent biological themes, and employs LLM-based summarisation to generate contextual interpretations of these themes.

GeneInsight uses a two-stage approach to extract and organise biological information from gene sets (Fig. 1a). In the biological theme generation stage, the system collects functional annotations from the STRING database for each input gene, creating a collection of gene-specific descriptions. This textual corpus is subjected to cluster-based topic modelling, which groups similar annotations into clusters (topics) and identifies key terms for each cluster. An LLM then converts representative annotations from each cluster into interpretable biological themes. These biological themes are linked back to genes via their associated descriptions, allowing selection of important themes via statistical testing (hypergeometric testing with false discovery rate correction) to identify which biological concepts are significantly enriched within the original gene set.

**Figure 1.**
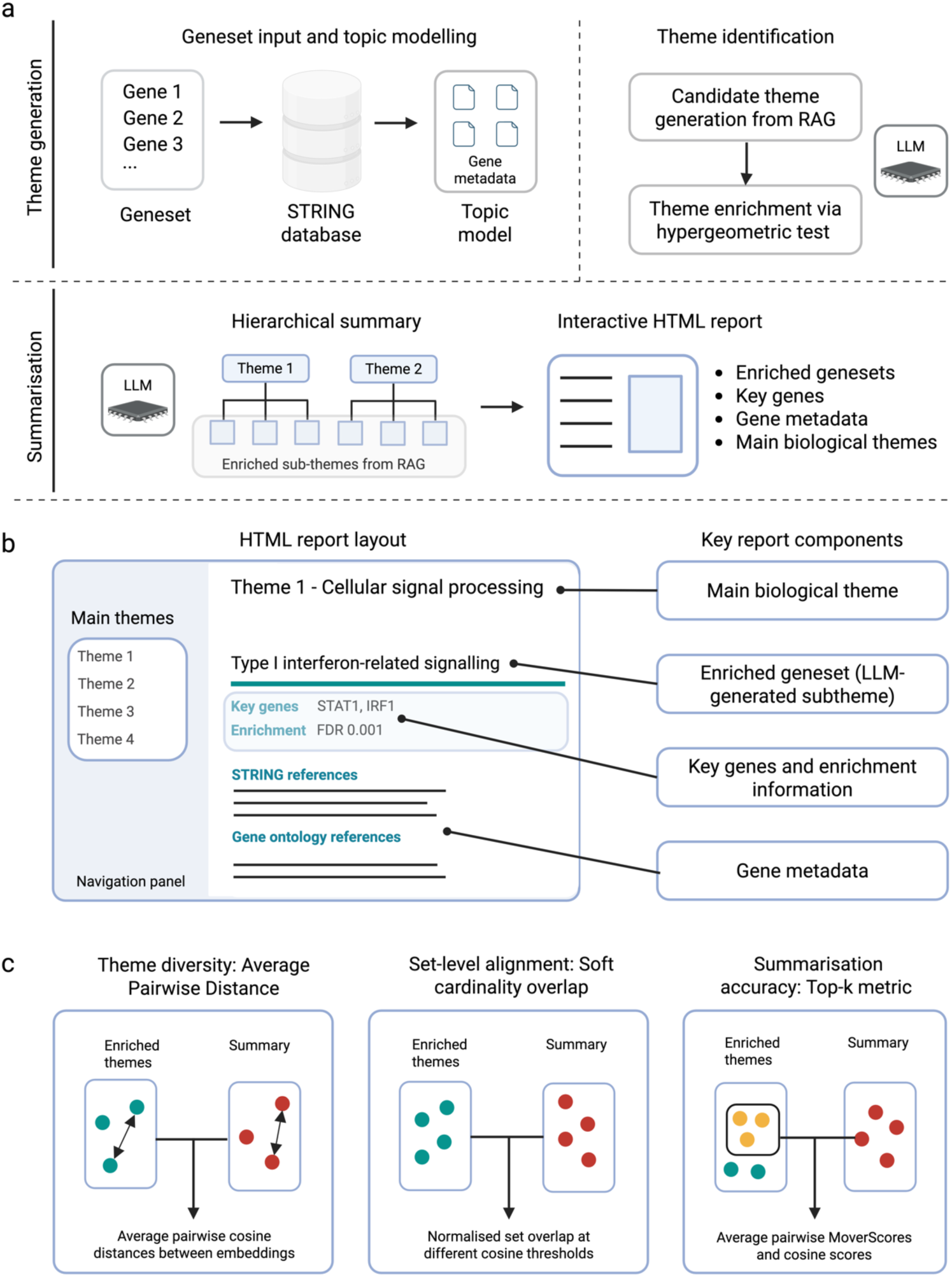
GeneInsight system architecture and workflow. (a) Schematic overview of the GeneInsight framework. The pipeline processes gene sets through two sequential stages: (1) Theme generation and (2) summarisation generates comprehensive reports with interactive visualisations. (b) Web interface components with components referenced in (a). (c) Performance metrics used in benchmarking showing theme diversity using Average Pairwise Distance, set-level alignment using Soft Cardinality Overlap and summarisation accuracy using Top-k metric. RAG, Retrieval Augmented Generation; LLM, Large Language Model; HTML, Hypertext Markup Language.

The second summarisation stage begins with another round of cluster-based topic modelling to identify key themes. This approach refines these enriched themes by measuring how consistently they appear as cluster representatives across multiple runs of topic modelling. The software then extracts the final summary by selecting themes to include based on user-defined length preferences. A large language model creates a hierarchical summary where major biological themes appear as main headings with related subheadings grouped beneath them. The final interactive HTML report (Fig. 1b) seamlessly links theme descriptions to their corresponding gene annotations. This integration enables researchers to easily navigate between overarching biological processes and their specific components. Moreover, every theme is directly tied to the original enrichment results, ensuring that all findings presented by GeneInsight are firmly grounded in rigorous statistical analysis.

We first characterised GeneInsight’s enrichment stage by comparing it with the STRING database functional enrichment application programming interface (API). Using identical underlying gene-level information and 1,000 Molecular Signatures Database (MSigDB)^9^ gene sets, this comparison directly assessed our tool’s effectiveness in identifying important biological themes. We measured performance through metrics (Fig. 1c) that evaluated both the diversity of identified concepts and the degree of overlap between methods.

GeneInsight consistently identified a larger number of enriched gene sets than the STRING database functional enrichment API across all significance thresholds (Fig. 2a, Supplementary Fig. 1). While the two methods show strong positive correlation (r=0.69-0.87), GeneInsight typically returns 2-3 times more enriched terms than the STRING-DB API at equivalent statistical thresholds (FDR<0.05). Notably, this pattern persists even at the most stringent significance levels (FDR<0.001) (Supplementary Fig. 1), suggesting that our gene-specific summarisation approach captures biological relationships complementary to those identified by STRING-DB’s enrichment method (Fig. 2b).

**Figure 2.**
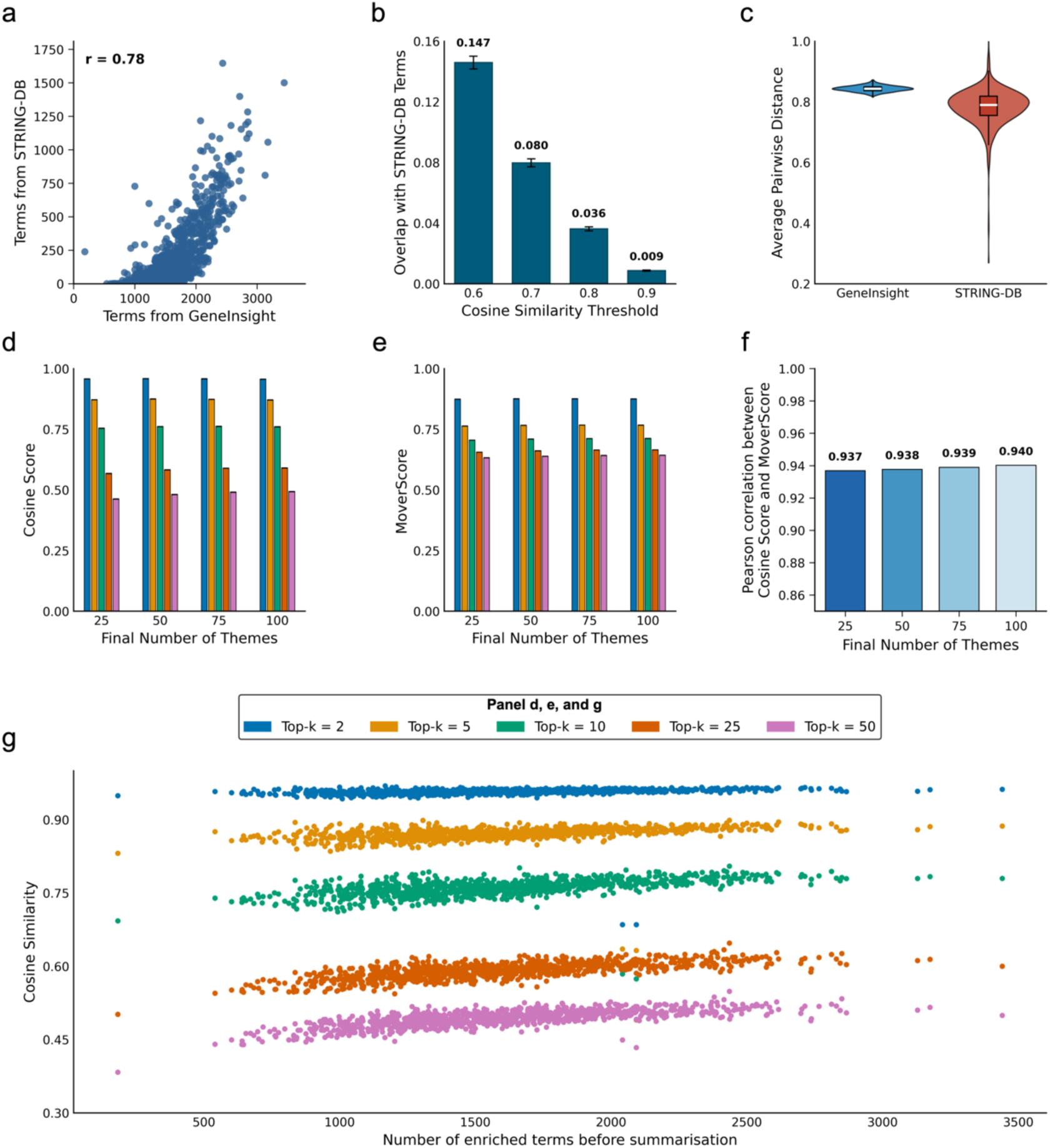
Evaluation of GeneInsight and STRING-DB. (a) Scatter plot showing the number of terms identified by GeneInsight versus STRING-DB API across 1,000 MSigDB gene sets. (b) Overlap of significant terms between GeneInsight and STRING-DB at various semantic similarity thresholds. (c) Average Pairwise Distance (APD) measurements comparing GeneInsight and STRING-DB enrichment results. (d) Top-k similarity scores for enriched gene sets across different user-defined summary levels (25, 50, 75, and 100 terms). (e) Top-k scores calculated using MoverScore for semantic similarity assessment. (f) Pearson correlation coefficients between cosine similarity and MoverScores at different levels of user-defined summarisation. (g) Top-k cosine similarity scores plotted against varying input corpus sizes (ranging from <500 to >3000 terms).

To evaluate the conceptual breadth of identified terms, we measured semantic diversity using average pairwise distance between terms. This metric quantifies how conceptually distinct each term is from all others in the set, with higher values indicating coverage of a broader range of biological concepts rather than redundant or closely related processes. GeneInsight-derived terms demonstrated significantly greater semantic diversity compared to STRING-DB terms (Fig. 2c), indicating its ability to capture a wider spectrum of biological information.

Next, we assessed the summarisation stage of GeneInsight going from these enriched themes to user-defined summaries. To evaluate theme preservation during summarisation at different levels (25, 50, 75, and 100 terms), we used a Top-k semantic similarity metric that focuses on each source term’s strongest matches to measure how well summaries capture essential biological concepts without being diluted by less relevant relationships. This approach identifies the k-nearest semantic neighbours for each term, better handling the imbalance between comprehensive source material and length-constrained summaries. GeneInsight demonstrated robust summarisation of key themes across different final summary counts (Fig. 2d), with the stability of these metrics suggesting that our tool effectively identifies core biological concepts regardless of user-supplied summary length constraints.

To provide an orthogonal validation beyond the cosine similarity metric, which measures the directional similarity between text representations, we employed Earth Mover’s Distance^11^ (MoverScore) (Fig. 2e). Like cosine distance, this complementary approach assesses semantic similarity by measuring the minimum cost required to transform one text into another in the embedding space, capturing different aspects of semantic relationships. All Top-k values demonstrated MoverScores greater than 0.5, indicating that each extracted theme successfully captured substantial semantic content from the original enriched gene sets. The results showed consistently high correlation values (Fig. 2f) between MoverScores and cosine similarity scores (0.93 - 0.94) across all theme configurations, confirming the robustness of our semantic similarity assessments.

We also confirmed that summarisation performance remained stable across documents of varying sizes (from <500 to >3000 terms), demonstrating that our tool’s summarisation capability is independent of input size (Fig. 2g).

Next, we applied GeneInsight to an RNA-seq dataset from a murine model of mesothelioma treated with immunotherapy, specifically focusing on genes differentially expressed in treatment responders^12^. GeneInsight prioritised biological themes centred around Type I interferon signalling and monocyte-macrophage axis activation (Supplementary Data 1) which were not detected through standard Gene Ontology (GO) enrichment analysis. The importance of Type I interferon signalling was subsequently validated through mouse models using antibody-mediated interferon blockade experiments, which confirmed the functional relevance of these pathways in treatment response. The monocyte involvement themes identified by our tool were further validated through single-cell analysis, confirming GeneInsight’s capacity to identify biologically relevant signatures that effectively bridge bulk and single-cell approaches.

We then evaluated GeneInsight on a multi-omics dataset from the DREAM^13,14^ study, which included mesothelioma patients undergoing chemoimmunotherapy treatment. Using responder-specific genes identified through a NanoString panel and bulk RNA-seq time course data, GeneInsight successfully extracted stem cell-like signatures in T-cells (Supplementary Data 2) that were independently confirmed through orthogonal single-cell validation. In the original analysis, these stem-like signatures were only discovered after weeks of analysis involving differential abundance testing, manual inspection of CD8^+^ T cell subclusters, differential expression analysis and marker analysis comparing responders and non-responder populations^13^. GeneInsight streamlined this process by automatically identifying these key biological themes in a single analysis in of 30 minutes, demonstrating how it can reduce analytical complexity and accelerate hypothesis generation from multi-omic datasets.

Finally, we evaluated GeneInsight using a published gene set^15^ associated with the transcriptional response of neutrophils to Francisella tularensis infection. GeneInsight identified distinct metabolic reprogramming signatures from differentially expressed genes involved in glucose metabolism (Supplementary Data 3), suggesting a metabolic shift that aligns with neutrophil functional alterations during infection. This metabolic reprogramming pattern, particularly involving key glycolytic regulators such as PFKL, was validated in a follow-up study^16^ using the same dataset 9 years later. This case demonstrates GeneInsight’s capacity to derive novel biological insights from existing datasets, potentially accelerating discovery timelines from functional genomics data.

## Discussion

Interpreting high-throughput genomic data remains a significant challenge as biological databases and literature continue to grow. GeneInsight addresses this by integrating large language models with topic modelling, capturing biological insights beyond what enrichment analysis offers. Furthermore, GeneInsight significantly reduces the time-consuming and potentially error-prone manual curation process traditionally required when reconciling redundant outputs from multiple enrichment analyses, allowing researchers to rapidly extract clinically relevant and actionable biological insights from increasingly complex genomic datasets.

GeneInsight’s algorithm is designed to process text-based annotations regardless of their source. While our current implementation uses the STRING database, the framework can readily incorporate PubMed abstracts, Gene Ontology terms, pathway descriptions from Reactome^17^, and unpublished private datasets. This extensibility allows GeneInsight to evolve alongside emerging knowledge sources while maintaining its core analytical capabilities. This approach represents a potential shift from gene set analysis using conventional, expert-defined ontologies to dynamic, LLM-powered systems that derive biological themes directly from expanding scientific literature.

## Supplementary data and figures

**Supplementary Figure 1.**
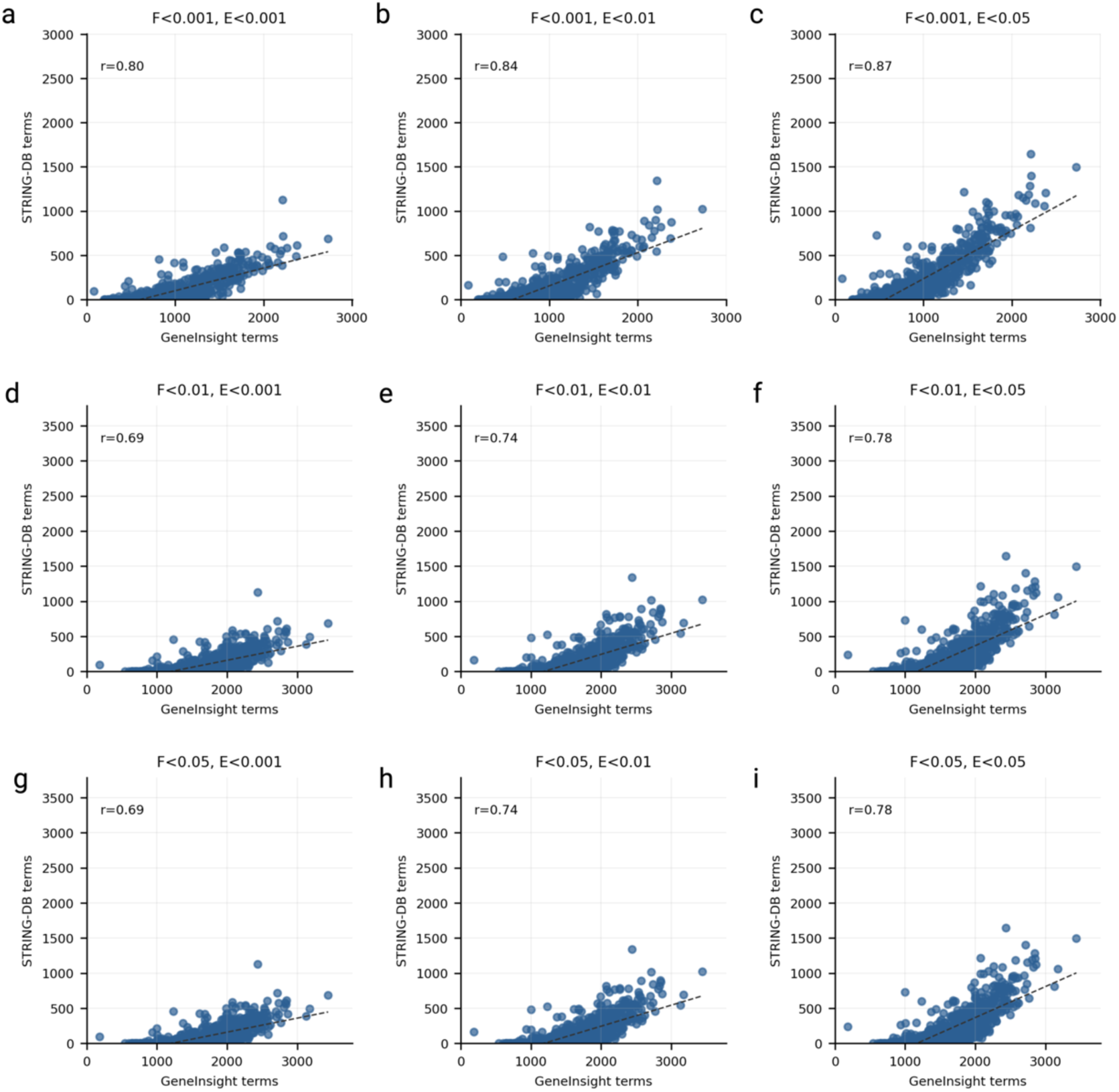
Correlation between GeneInsight and STRING database (STRING-DB) enriched gene sets at different significance thresholds. The grid shows pairwise comparisons of enriched gene sets identified by GeneInsight (x-axis) versus terms from the STRING database (y-axis) at varying statistical significance thresholds. Each subplot represents a different combination of p-value thresholds, with F indicating the adjusted p-value threshold (FDR) for GeneInsight results and E indicating the FDR threshold for STRING database results. Thresholds range from stringent (0.001) to permissive (0.05). Blue dots represent individual gene sets with the number of enriched terms plotted for each tool. Dashed lines show the linear regression trend. Correlation coefficients (r) are displayed in the upper left of each panel.

**Supplementary Figure 2.**
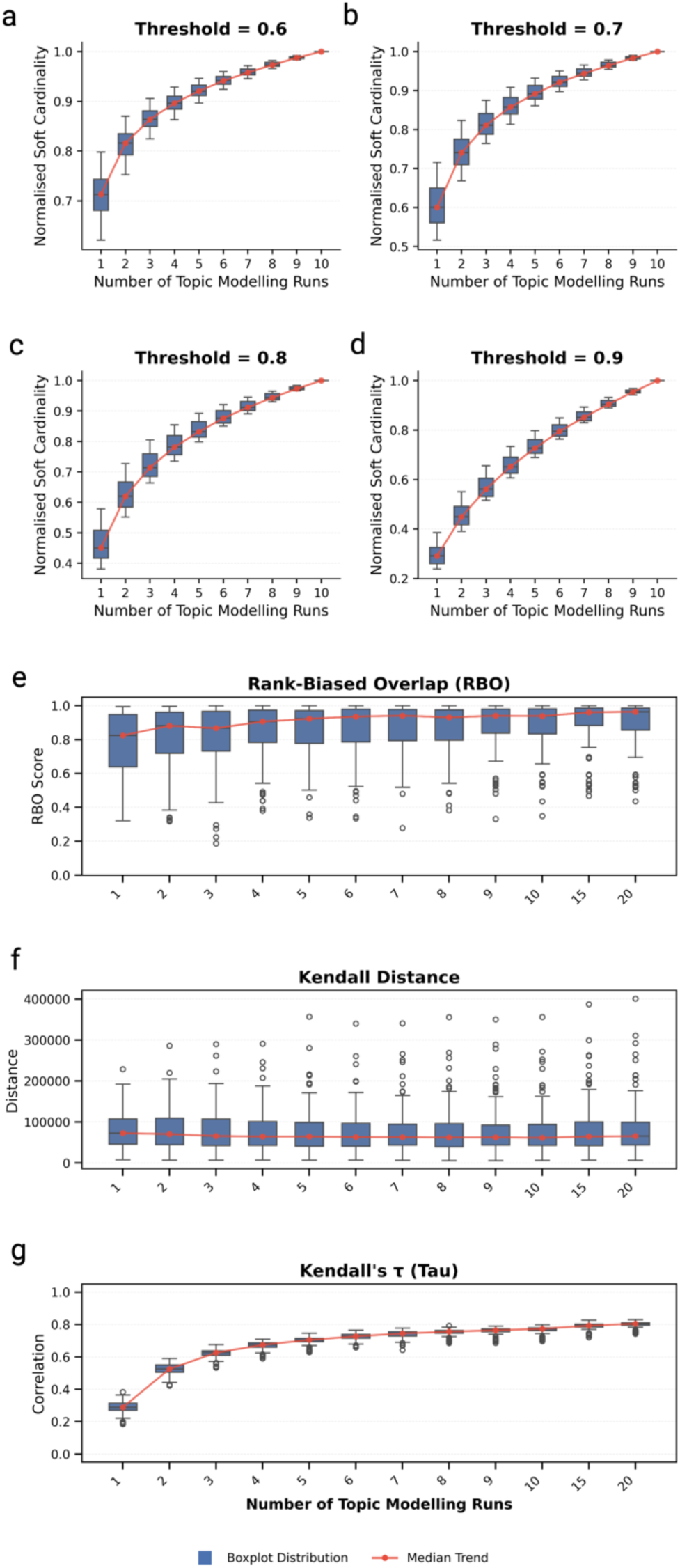
Convergence analysis of topic modelling stability with increasing number of runs. Top panels (a-d): Normalised soft cardinality measures the semantic overlap between terms identified in each individual topic modelling run and the complete term set across all runs. Higher values indicate greater coverage of the overall vocabulary space. Bottom panels (e-g): Box plots show three complementary ranking metrics comparing each run (1-20) against the reference (25 runs): Panel e (Rank-Biased Overlap, RBO), Panel f (Kendall Distance), and Panel g (Kendall’s τ correlation). Higher RBO and Kendall’s τ values indicate greater similarity to reference rankings, while lower Kendall Distance values represent better alignment. For all panels, trend lines connect median values, showing the stabilization pattern as the number of topic modelling runs increases. Box plots show distribution (centre line = median, box limits = first and third quartiles, whiskers = 1.5× interquartile range, points = outliers).

**Supplementary Data 1.** HTML reports and raw files from a GeneInsight analysis of the murine mesothelioma immunotherapy response dataset, highlighting biological themes related to Type I interferons and monocyte-macrophage activation.

**Supplementary Data 2.** HTML reports and raw files from a GeneInsight analysis of the DREAM study mesothelioma patient dataset, highlighting stem cell-like (lymphocyte proliferation and differentiation) signatures in T-cells of responders to chemoimmunotherapy treatment.

**Supplementary Data 3.** HTML reports and raw files from a GeneInsight analysis of neutrophil transcriptional response to Francisella tularensis infection, featuring identified metabolic reprogramming signatures in glucose metabolism.

## Online methods

### Implementation and benchmarking workflow

GeneInsight is implemented in Python (3.10+). The workflow management system Snakemake (v7.32.4) coordinates the benchmarking pipeline, while documentation is generated using Sphinx (v7.2.6). Core computational components include BERTopic (v0.15.0) for topic modelling and Optuna (v3.3.0) for hyperparameter optimisation.

### Source data

The STRING database serves as a fundamental data source for GeneInsight, providing comprehensive protein-protein interaction (PPI) data across multiple organisms. We utilised STRING version 12.0. The database integrates both direct physical interactions and indirect functional associations, drawing from multiple evidence types including experimental data, pathway knowledge, co-expression patterns, and text mining of scientific literature.

For our analysis pipeline, we specifically leverage STRING’s functional enrichment API, which provides statistically validated functional annotations for protein sets. The API accepts protein identifiers and returns enriched terms across multiple functional classification systems, including Gene Ontology (GO) terms, KEGG pathways, and UniProt^18^ keywords. Each returned term is accompanied by a false discovery rate (FDR), p-value, and the number of proteins from the query set that map to that term.

### Algorithm description

GeneInsight employs a two-stage approach for gene set analysis that processes biological information at the individual gene level before performing knowledge synthesis, offering an alternative to conventional set-level enrichment methods.

In the first stage (biological theme generation), we query the STRING functional enrichment API to retrieve associated terms for each gene in the input set based on their statistical significance. This gene-level querying, as opposed to direct set-level enrichment, allows us to capture the full spectrum of functional annotations per gene, forming a rich document corpus that preserves gene-specific context.

BERTopic (v0.15.0) then processes this corpus through cluster-based topic modelling to identify coherent biological themes by grouping similar annotations into clusters. Each topic represents a collection of functionally related terms supported by gene-specific literature evidence. A large language model (GPT-4o mini by default) converts these detailed topic representations into interpretable biological themes while maintaining the granular insights gained from per-gene analysis.

To link each theme back to genes, each LLM-generated topic serves as a search query against the corpus. Since each document maintains its link to specific genes, we create gene sets for each topic by combining genes from the top matching documents (default N=5). We then validate these topic-associated gene sets using hypergeometric testing against user-provided query genes and background sets (with Benjamini-Hochberg correction for false discovery rate), identifying which biological concepts are significantly enriched within the original gene set.

The second stage (summarisation phase) begins with another round of cluster-based topic modelling to identify key themes. Our approach ensures robust theme identification by performing multiple independent rounds (default N=5) of topic modelling with different random seeds. The terminal condensation phase identifies stable, biologically meaningful topics that persist across initialisations by measuring how consistently they appear as cluster representatives. We identify these stable topics through cosine similarity measurements between iterations and rank them by frequency.

The final topic selection uses similarity thresholds that align with user-specified enrichment criteria and length preferences, facilitated by automatic optimisation against clustering metrics. Specifically, we employ Davies-Bouldin^19^ and Calinski-Harabasz^20^ scores to evaluate cluster quality, which quantify both the cohesion within clusters and the separation between them. The LLM then creates a hierarchical summary where major biological themes appear as main headings with related subheadings grouped beneath them, presented in an interactive HTML report that connects theme descriptions directly to the original gene annotations, allowing researchers to easily navigate between broad biological processes and their specific components while maintaining accuracy to the underlying gene functions.

### Semantic Overlap Metric for Biological Term Sets

We developed a semantic overlap metric to quantify relationships between biological term sets, defined as SC(A,B) = Σ max(cos(e_a_,e_b_)) for all a ∈ A, b ∈ B, where e_a_ and e_b_ represent embedding vectors of terms from sets A and B respectively, and cos denotes cosine similarity. This metric allows semantically similar phrases to be counted as overlapping, rather than requiring exact matches. To enable comparison across datasets of varying sizes, we implemented a normalised form NSC(A,B) = SC(A,B) / |A|, where |A| represents the cardinality of set A. Dense vector representations were generated using the SentenceTransformer framework with paraphrase-MiniLM-L6-v2, producing 384-dimensional embeddings that capture contextual relationships between terms.

We applied this metric in two contexts: first, to optimize BERTopic hyperparameters by comparing different numbers of sampling rounds during the first phase (n = 1 to n = 10) and assessing the stability of the resulting term sets; and second, to evaluate the overlap between GeneInsight-derived themes and conventional STRING-DB functional enrichment results, providing quantitative measures of concordance between these different approaches (Fig. 2b).

### Top-k Semantic Similarity Analysis

Assessing theme preservation in biological term sets requires robust quantification of meaningful relationships while accounting for the inherent variability in gene set annotations. To evaluate how well summaries preserve core themes from source terms, we developed a recall-based similarity metric that captures nuanced relationships beyond exact-match approaches. This metric specifically measures how well the summary captures essential ideas from the source material, rather than selective details. Low recall would indicate missing themes, while high recall ensures comprehensive preservation of key biological concepts.

For any two sets of terms A = {a₁, a₂,…, aₙ} and B = {b₁, b₂,…, bₘ}, we compute dense vector representations using the all-MiniLM-L6-v2 model, generating embedding matrices E_A_ ∈ ℝⁿˣᵈ and E_B_ ∈ ℝᵐˣᵈ, where d = 384 represents the embedding dimension. For each term i in set A, we compute its k highest similarity scores with terms in set B using cosine similarity: 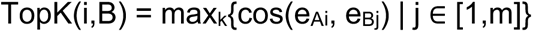. The overall similarity score is calculated as 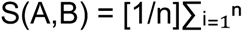 mean(TopK(i,B)), where the mean averages the k highest similarities for each term.

We evaluate different k values (k = 2, 5, 10, 25, 50) to provide complementary views of how well meaning is retained between term sets. Lower k values focus on the strongest conceptual matches, while higher values assess broader thematic coverage. This multi-k assessment strategy reveals whether high similarity scores are maintained across different matching thresholds.

The Top-k approach offers several advantages over simple average similarity. By focusing on each source term’s strongest matches rather than averaging over all pairwise comparisons, the method prevents dilution of the most meaningful relationships and ensures that high similarity scores from key themes drive the overall metric. This approach improves robustness to noise and outliers by considering only the TopK similarities for each term, making the metric less sensitive to low-scoring, less relevant matches. The method is particularly effective for handling imbalanced sets, where many terms in a larger set may not have close equivalents in the summary. By focusing solely on the strongest conceptual relationships, the approach provides a clearer assessment of how well the summary captures essential content.

The multi-threshold analysis using different k values provides insights into both the strongest matches and how comprehensively the final HTML summary retains information across user-defined length constraints. We specifically evaluated summaries containing between 25 and 100 terms, assessing how effectively these concise summaries maintained the core biological concepts from the more extensive source material. This approach is particularly advantageous when evaluating how well a summary preserves core themes from a larger set, as it emphasises conceptually significant matches, reduces noise from irrelevant pairings, and effectively handles set size imbalances inherent in summarisation tasks with strict length limitations.

### Hyperparameter Optimisation

We utilised the Molecular Signatures Database (MSigDB) C2 collection as our primary data source for both validation and testing. This collection provides a comprehensive resource of curated gene sets derived from pathway databases, published experiments, and expert knowledge. Each gene set includes detailed text descriptions of its experimental derivation, offering both transcriptomic and semantic information. We randomly selected 100 gene sets for validation and 1000 distinct gene sets for testing, ensuring comprehensive coverage across diverse biological processes and experimental conditions.

Hyperparameter optimisation on the validation set focused on two critical aspects of GeneInsight’s implementation: the biological theme identification (theme generation phase) parameters (Supplementary Figure 2a-d) and the rank-based clustering iterations (summarisation phase) (Supplementary Figure 2e-g). For the biological theme identification phase, we employed an elbow method using soft cardinality intersection against a term universe generated through exhaustive sampling across 10 initialisation seeds across 1-10 sampling runs. For the summarisation phase, we evaluated convergence using rank-based metrics with an upper bound of 100 biological theme identification runs. Through systematic evaluation of cosine similarity thresholds and soft overlap intersection measures, we determined optimal cut-off points based on clustering stability analysis. This process led to default values of 5 sampling rounds and 10 rank sampling rounds, selected for their empirical performance on the validation set.

### Comparative evaluation

For comparative evaluation against conventional approaches, we assessed GeneInsight’s performance against STRING-DB functional enrichment using an independent test set of 1000 MSigDB gene sets. Enrichment analysis was performed using STRING-DB (version 12.0). For each gene set, we computed three complementary metrics to evaluate term diversity and semantic preservation.

Term diversity was quantified using Average Pairwise Distance (APD) between semantic embeddings. For each term set, we generated dense vector representations using the all-MiniLM-L6-v2 model (384-dimensional embeddings) and computed pairwise cosine distances. The APD was calculated as the mean of these distances, with higher values indicating greater semantic diversity. Statistical significance of the differences between GeneInsight and STRING-DB results was determined using a two-sided t-test.

Semantic preservation was assessed using the Top-k similarity metric (described above), where for each source term, we identified the k most semantically similar terms in the summarised set (k = 2, 5, 10, 25, 50) using cosine similarity between embedding vectors. As an orthogonal validation approach complementary to cosine similarity but not incorporated into the GeneInsight software itself, we also evaluated performance using the MoverScore, which measures optimal transport distance between term embeddings and accounts for both semantic and positional alignment.

Performance was evaluated across different user-specified biological theme counts in the final HTML report (25, 50, 75, and 100 themes) to assess how well the key biological concepts were maintained at various summary lengths.

### Dataset Analyses

The runs were performed using default parameters for topic modelling with a specified summary length of 100 terms and a temperature parameter of 0.0, followed by manual inspection of enriched terms and the HTML summary.

For the first dataset, we analysed a collection of "fast-off" genes as defined in a time-course analysis^12^ of the AB1 mesothelioma mouse model across four time points following immune checkpoint blockade administration.

In our second dataset, we examined RNA-seq data from mesothelioma patients containing time-course expression profiles, as described in the Phase II DREAM^13^ chemo-immunotherapy study. We specifically focused on the first 100 genes identified as important responders based on their factor loading scores. Analysis was performed using DIABLO^21^, which enabled the identification of key predictors from multi-omic data.

For the third dataset, we analysed a published gene set^15^ associated with the transcriptional response of neutrophils to Francisella tularensis infection (GSE37416). This gene set specifically comprised downregulated genes identified from human polymorphonuclear leukocytes (PMN) treated with F. tularensis subsp. holarctica live vaccine strain (LVS) for 48 hours compared to untreated controls at 0 hours.

## Notes

https://github.com/wlchin/geneinsight

https://wlchin.github.io/geneinsight/index.html

### Competing Interest Statement

The authors have declared no competing interest.

